# Transcriptional Signatures of Field Cancerization in Gastric Cancer

**DOI:** 10.1101/2025.09.15.676327

**Authors:** Jéssica Manoelli Costa da Silva, Ronald Matheus da Silva Mourão, Juliana Barreto Albuquerque Pinto, Diego Pereira, Valéria Cristiane Santos da Silva, Kauê Sant’Ana Pereira Guimarães, Ana Karyssa Mendes Anaissi, Samia Demachki, Williams Fernandes Barra, Samir Mansour Casseb, Fabiano Cordeiro Moreira, Rommel Mario Rodriguez Burbano, Paulo Pimentel de Assumpção

## Abstract

The high rate of local recurrence in gastric adenocarcinoma (GA) suggests that carcinogenesis is not a focal event but a field-wide process. This phenomenon, known as “field cancerization,” posits that histologically normal peritumoral tissue is, in fact, a pre-neoplastic field harboring incipient molecular alterations that confound genomic studies using it as a normal control. To overcome this limitation, we performed a three-way comparative transcriptomic analysis of tumor, peritumoral, and true-normal gastric tissues using a deep learning framework. We identified a stable 138-gene signature established within the peritumoral field and conserved in the tumor, which was absent in healthy controls. Within this signature, three key COSMIC-listed driver genes were highlighted: the Hippo pathway component *FAT4* and the p53-inhibitor *MDM4* were upregulated, while the EMT-suppressor *NDRG1* was repressed. Co-expression analysis revealed a dynamic rewiring of these drivers, with a significant positive correlation between *FAT4* and *MDM4* emerging exclusively in the peritumoral field. In contrast, a negative correlation between FAT4 and NDRG1 was observed specifically in the tumor context. In public cohorts, high expression of *FAT4* and *MDM4* was significantly associated with poor patient prognosis, whereas *NDRG1* showed no prognostic association. Critically, the prognostic power of *MDM4* was validated in our local patient cohort. Our findings demonstrate that the peritumoral field is a molecularly distinct state in gastric carcinogenesis, characterized by a metabolic shift, and identify *FAT4* and *MDM4* as key drivers of this early transition, with significant potential as prognostic biomarkers.

## Introduction

The molecular and clinical heterogeneity of gastric adenocarcinoma (GA) represents a major obstacle to improving patient outcomes, limiting the effectiveness of prognostic stratifications and therapeutic interventions. This intrinsic variability is not adequately captured by conventional histopathological classifications (Laurén, 1965). In response, significant oncology research has focused on developing robust molecular classifications, culminating in systems such as the TCGA, which are designed to align genomic profiles with clinical outcomes and therapeutic sensitivity.

However, a critical methodological factor permeates many of these studies: the use of adjacent peritumoral tissue as a normal biological control. The use of peritumoral tissue for this purpose can be problematic, as the “field of cancerization” theory (Slaughter et al., 1953) suggests that it already exists in a molecularly altered state. This implies that peritumoral tissue may share a subset of molecular aberrations with the tumor, such as mutations in driver genes, potentially obscuring the identification of those events that are truly specific to cancer initiation and progression (Curtius et al., 2018; de Assumpção et al., 2018).

The resolution of the transcriptomic programs that govern this malignant conversion requires high-throughput methodologies. While RNA-sequencing (RNA-seq) provides the necessary depth of transcriptomic sampling, the resulting high-dimensional datasets contain complex, non-linear structures that are intractable by conventional statistical methods. This necessitates the application of advanced computational models, such as deep learning, capable of modeling these complex interactions and identifying latent predictive signatures in the expression profiles (Mortazavi et al., 2008; Necula et al., 2019; Stark et al., 2019; Sun and Chen, 2023).

Therefore, this study was designed to address these challenges through a three-way comparative transcriptomic analysis of gastric adenocarcinoma, adjacent peritumoral, and control tissues from healthy donors. The inclusion of a normal control group is methodologically crucial, as it provides a biological baseline for distinguishing transcriptional signals. This allows for a rigorous distinction between changes attributable to the “field effect” and those intrinsically associated with the malignant phenotype.

## Materials and Methods

### Study Design and Data

We performed a comparative transcriptomic analysis using RNA-sequencing (RNA-seq) data from 72 paired gastric tumor and peritumoral tissues from our local cohort (Pará, Brazil; Ethics Approval CAAE: 47580121.9.0000.5634) and 68 normal gastric tissues from NCBI BioProject PRJNA1054173.

### RNA-Seq and Bioinformatic Processing

Total RNA was extracted from local samples (RNA Integrity Number > 7) and sequenced on a NextSeq platform. The bioinformatic pipeline included quality filtering (Fastp), transcript quantification (Salmon, Gencode v43), and subsequent gene-level normalization and variance stabilization (`DESeq2` rlog).

### Deep Learning and Feature Selection

A deep autoencoder with a 64-neuron bottleneck was trained in Keras/TensorFlow to learn a compressed representation of the gene expression data. The feature importance of each gene within the model’s latent space was then quantified using SHAP (SHapley Additive exPlanations), and the most relevant genes were selected via the Elbow method.

### Field Signature Identification and Statistical Analysis

We derived a carcinogenic-field signature by sequentially filtering genes: (i) SHAP-selected as relevant in both tumor and peritumoral tissues; (ii) stable between paired tumor–adjacent samples (Wilcoxon *p* > 0.9); (iii) dysregulated versus normal (|log □ FC| > 1, FDR < 0.05); and (iv) listed in COSMIC. Co-expression was summarized by Spearman correlation. UMAP (R umap; Euclidean, *n_neighbors* = 15, *min_dist* = 0.1) was computed on VST-normalized (DESeq2) expression restricted to the 138 genes. Per-sample signature activity was quantified with ssGSEA (GSVA) on the same matrix, yielding one enrichment score per sample. Group differences (normal, adjacent, tumor) were assessed with Kruskal–Wallis followed by pairwise Wilcoxon tests with Benjamini–Hochberg correction.

### Prognostic and Survival Analysis

The clinical relevance of the identified driver genes was assessed via a two-tiered survival analysis. First, the association between gene expression and overall survival (OS) was investigated in public gastric cancer cohorts using the Kaplan-Meier Plotter online database. For internal validation, survival analysis was performed on a subset of 46 patients from our Pará cohort with available clinical follow-up data. Kaplan-Meier curves were generated using the `survminer` and `survival` R packages, and differences in OS were assessed using the log-rank test. A p-value < 0.05 was considered statistically significant.

## Results

### Training and Performance of Deep Autoencoder Models

To generate a low-dimensional representation of the gastric cancer transcriptome, we trained deep autoencoder models on mRNA and lncRNA expression profiles from tumor and peritumoral tissues. All models showed stable and efficient convergence over 800 epochs, with loss functions decreasing progressively to near-zero values, which confirmed robust training without significant overfitting (Fig. S1). We then evaluated model performance by quantifying the mean squared error (MSE) between the original and reconstructed expression profiles. For mRNA profiles, the reconstruction error was substantially higher in tumor samples (MSE = 2.71 x 10 □□) compared to their peritumoral counterparts (MSE = 9.55 x 10 □□), reflecting greater transcriptomic complexity. In contrast, for lncRNA profiles, the difference was minimal between tumor (MSE = 7.99 x 10 □□) and peritumoral (MSE = 7.14 x 10 □□) tissues. The overall low magnitude of these errors underscores the models’ capacity to effectively capture the latent structure of the data, validating their use for downstream analysis (Fig. S2). □

#### Deep Learning Identifies a Shared Transcriptomic Landscape Between Tumor and Peritumoral Tissues

To define the gene expression programs active in gastric carcinogenesis, we first applied a deep autoencoder to the transcriptomes of tumor and peritumoral tissues, followed by SHAP analysis to determine the importance of each gene. This approach identified 10,609 mRNAs as relevant in the tumor context, and 9,483 mRNAs in the peritumoral context.

A comparative analysis revealed a substantial overlap between the two tissue types. We found that 9,009 mRNAs were shared, providing strong evidence that the peritumoral field possesses a complex molecular architecture similar to the established tumor (Fig. 1A).

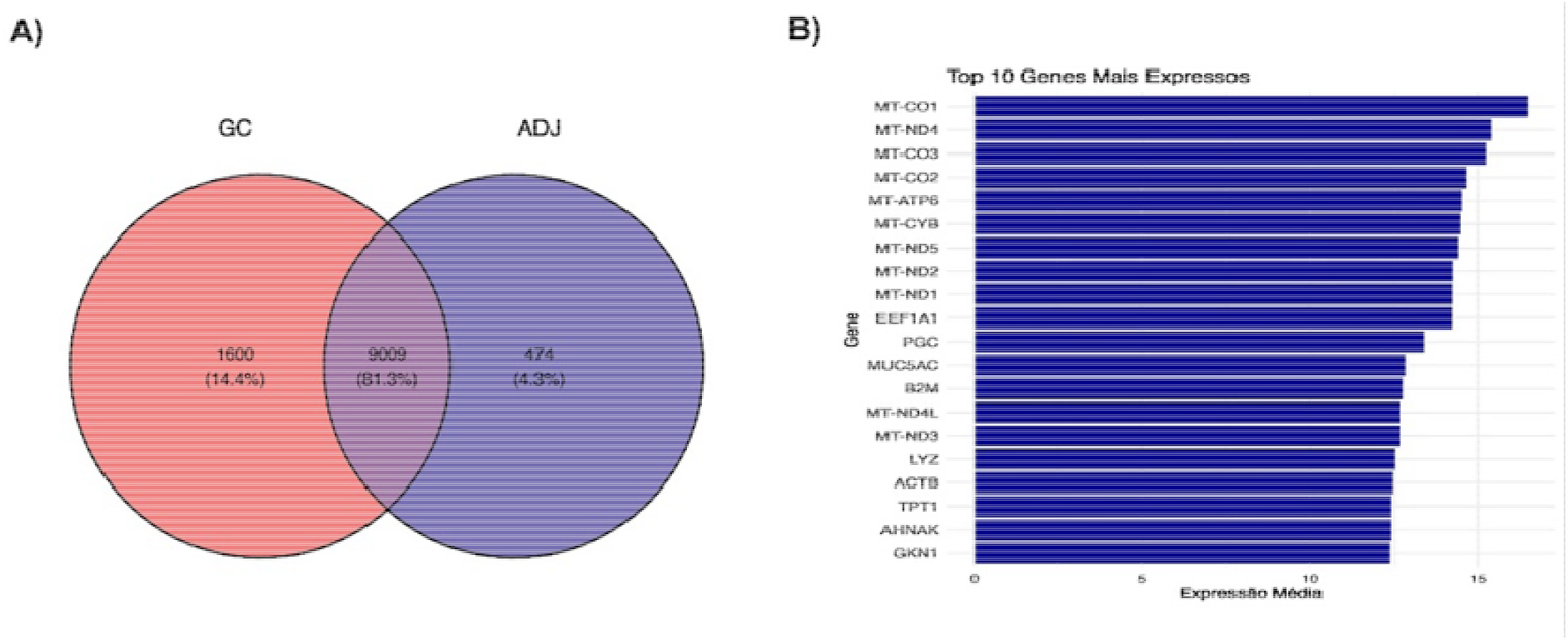

Analysis of the most highly expressed shared transcripts highlighted a mixture of both physiological and pro-oncogenic genes. Among the top shared mRNAs were genes essential for normal gastric function, such as those involved in mitochondrial respiration and epithelial protection (Fig. 1B).

#### A Stringent Statistical Pipeline Isolates a Core 138-Gene Carcinogenic Field Signature

Starting from the 9,009 genes shared between tumor and peritumoral tissues, we aimed to isolate a core program specific to the carcinogenic field. We applied a conservative stability filter, retaining only genes with indistinguishable expression in paired tumor–peritumoral samples (Wilcoxon test, p > 0.9). This reduced the list to a 138-gene subset that captures alterations already established in the peritumoral compartment and preserved in the tumor.

To verify that this signature is absent from healthy mucosa, we tested the 138 genes against the normal tissue cohort. All 138 were significantly dysregulated relative to normal (|log□FC| > 1, FDR < 0.05), within the broader set of 9,647 differentially expressed genes detected in the tumor-vs-normal contrast (Fig. 2A). A Venn diagram summarizes the selection workflow (Fig. 2B). Panel 2C highlights the expression patterns of the top 10 representative genes— *ADAMTS1, ITGB3, MST1R, NDRG1, MDM4, IRS2, ACSS1, MSMO1, FAT4*, and *XBP1*—across normal, peritumoral, and tumor tissues, illustrating a consistent gradient of dysregulation aligned with the carcinogenic process.

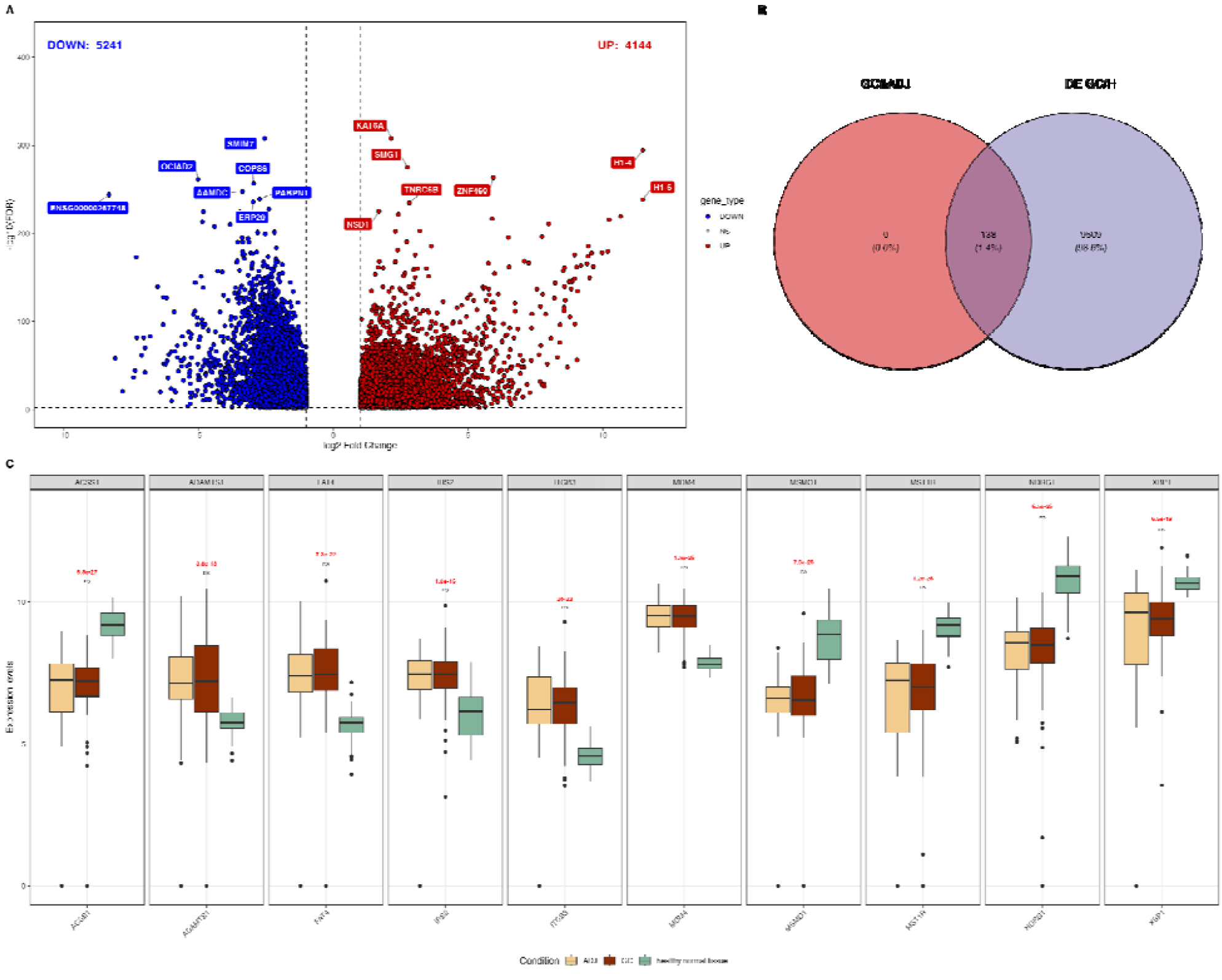

Collectively, these analyses define a 138-gene transcriptomic signature of the gastric carcinogenic field—present and stable in peritumoral tissue, maintained in tumor, and distinctly absent in normal mucosa—which we subsequently used for driver prioritization and functional interrogation.

#### The 138-Gene Signature Segregates the Carcinogenic Field from Normal Tissue

Using the 138-gene set, Uniform Manifold Approximation and Projection (UMAP) embedding revealed a clear separation between healthy gastric mucosa and diseased samples (Fig. 3A). Normal tissues aggregated into a compact, isolated cluster, whereas adjacent non-tumoral (ADJ) and gastric cancer (GC) samples colocalized within the same manifold, with partial internal separation. This configuration indicates that ADJ and GC share a common transcriptional program that is already established in the peritumoral field and is distinct from the normal state.

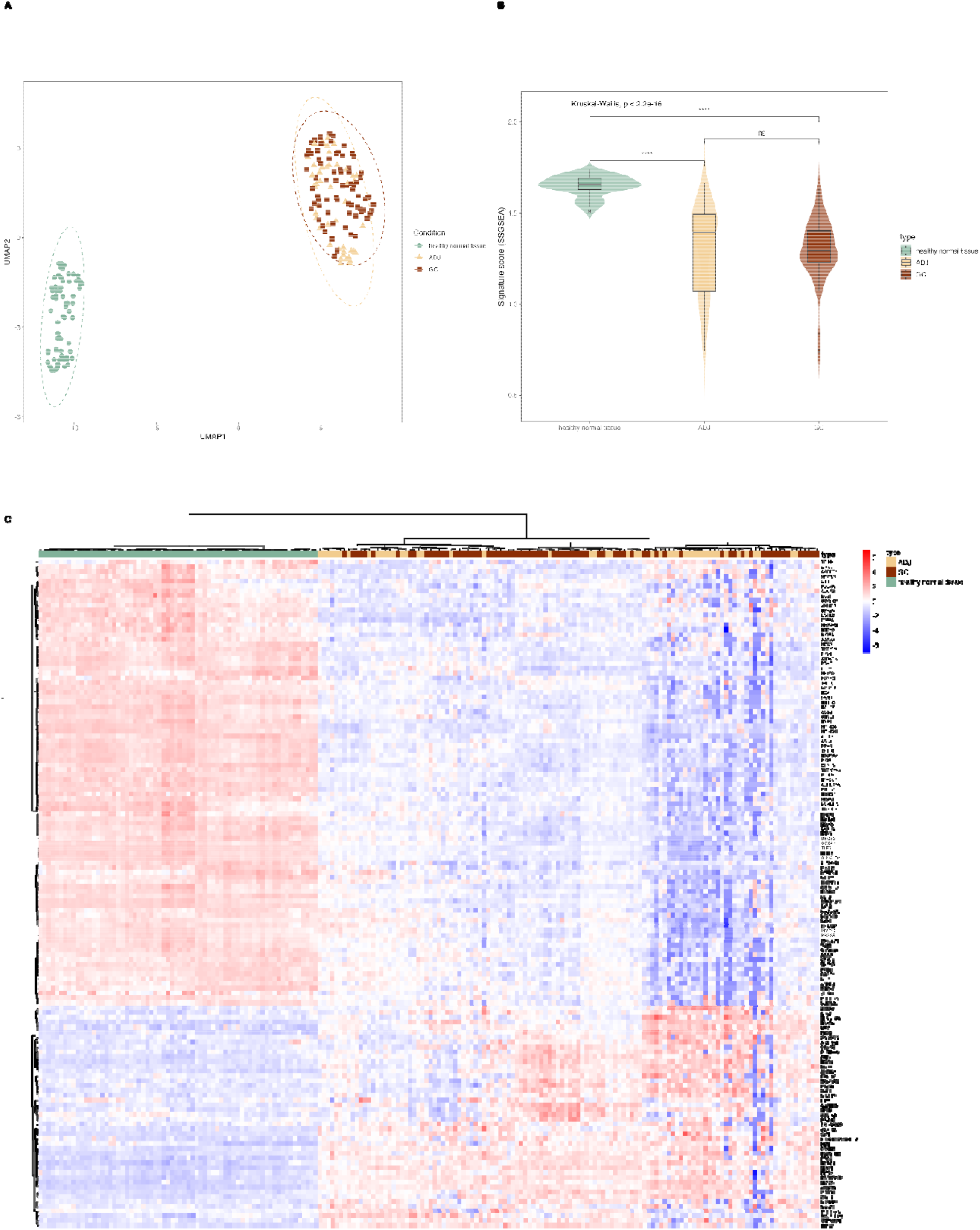

We next quantified the signature at the sample level using single-sample GSEA (ssGSEA). Signature enrichment scores differed significantly among the three groups (Kruskal–Wallis, p< 2.2 × 10□^1^□; Fig. 3B). Post-hoc pairwise tests showed higher scores in ADJ and GC relative to normal (both significant after multiple-testing adjustment), whereas the ADJ–GC contrast was not significant, indicating that the field program is activated in peritumoral tissue and sustained in tumor.

Unsupervised hierarchical clustering based on the same gene set reproduced this structure (Fig. 3C), yielding two primary sample clades: one comprising normal tissues and another containing ADJ/GC, which were largely intermingled. The heatmap displayed a graded pattern (normal → ADJ → GC) and resolved three co-expression modules: two modules upregulated in the carcinogenic field (ADJ/GC) relative to normal, and one module downregulated in the field. Together, these analyses demonstrate that the 138-gene set captures a coherent transcriptomic program of field cancerization—present in peritumoral tissue, maintained in tumor, and absent from normal mucosa.

#### Core Driver Genes of the Field Signature Display Dynamic Regulatory Rewiring

To prioritize the most functionally significant genes within the 138-gene signature, we cross-referenced the list with the COSMIC Cancer Gene Census, which identified three established cancer driver genes. The Hippo pathway component *FAT4* (log □ FC = +2.00) and the p53-negative regulator *MDM4* (log □ FC = +1.71) were both significantly upregulated, while the putative metastasis suppressor *NDRG1* was repressed (log □ FC = −2.00).

To understand how these drivers operate within the carcinogenic field, we next investigated their regulatory dynamics in both peritumoral and tumor contexts. This revealed a striking, context-dependent rewiring of their relationships (Fig. 4). In the peritumoral field, a significant positive correlation emerged between *FAT4* and *MDM4* (Spearman’s ρ = +0.35). Conversely, in the tumor context, this association was lost, and a distinct negative correlation appeared between *FAT4* and *NDRG1* (ρ = −0.36).

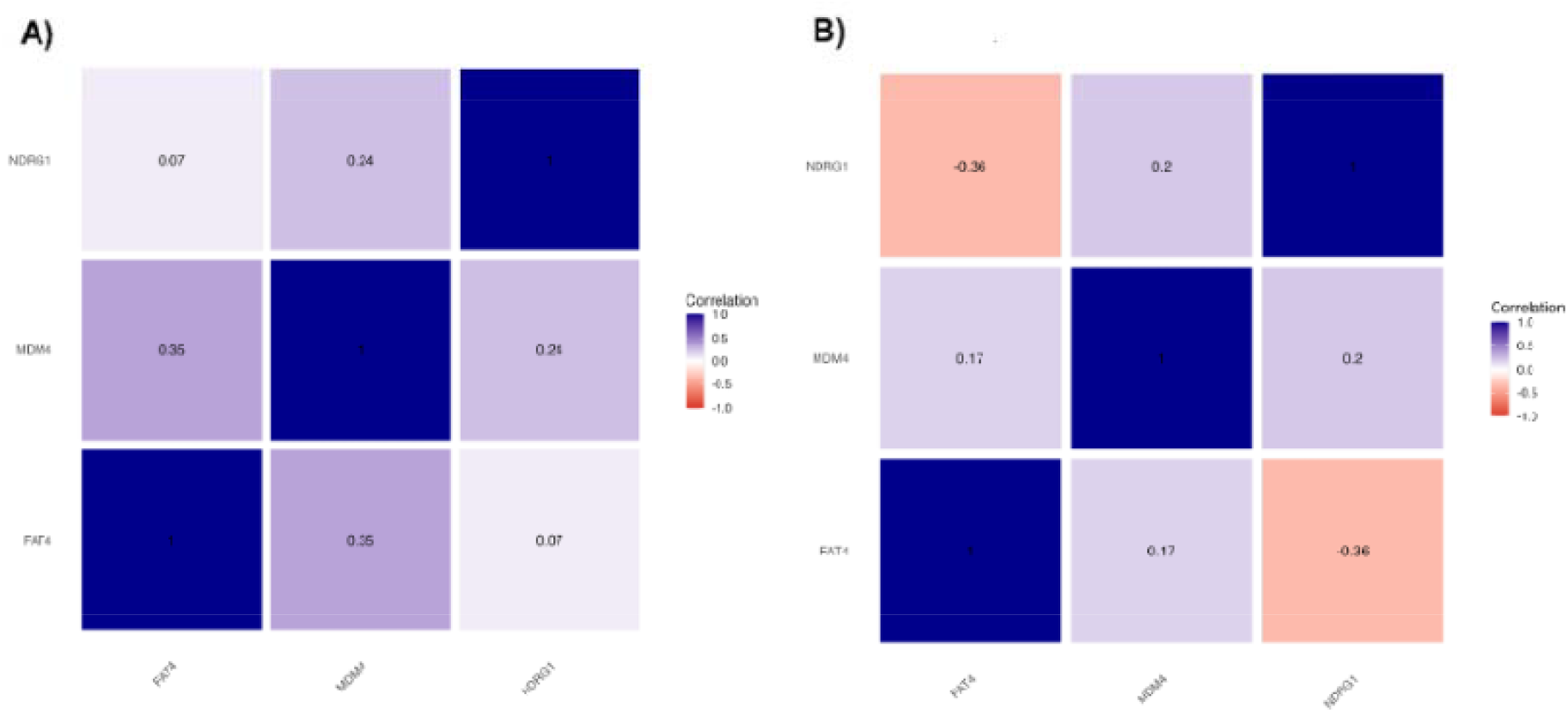

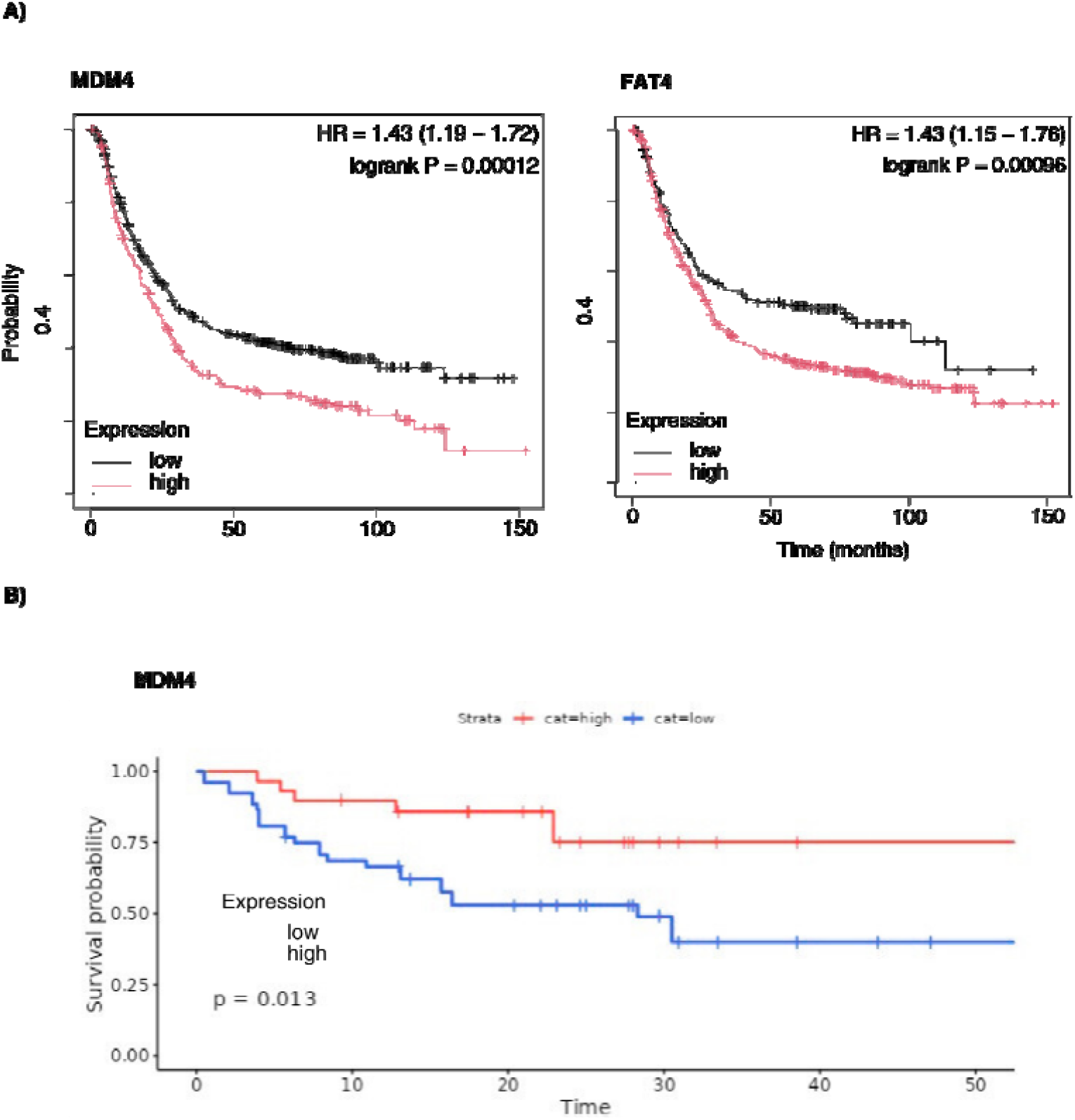

Together, these results not only identify the key drivers established in the gastric carcinogenic field but also reveal that their regulatory network is dynamically rewired during malignant progression.

#### High Expression of *FAT4* and *MDM4* is Associated with Poor Prognosis in Gastric Cancer

To ascertain the clinical relevance of the three identified driver genes, we evaluated their prognostic value. The analysis demonstrated that high expression of both *FAT4* and *MDM4* was significantly associated with worse overall survival. Patients with high levels of *FAT4* had a Hazard Ratio (HR) of 1.43 (95% CI: 1.15 – 1.76), with a log-rank p-value of 0.00096. Similarly, high expression of *MDM4* was linked to a poor prognosis, showing an identical HR of 1.43 (95% CI: 1.19 – 1.72) and a log-rank p-value of 0.00012. In contrast, the expression of *NDRG1* showed no significant association with patient survival (HR = 0.92 [95% CI: 0.78 – 1.09]; log-rank P = 0.35). These findings from large-scale public cohorts establish the prognostic potential of *FAT4* and *MDM4* as markers of adverse outcomes in gastric cancer, prompting their validation in our local patient cohort.

## Discussion

The present study confronts two of the most critical challenges in gastric oncology: the deep molecular heterogeneity of the disease and the methodological constraint of using peritumoral tissue as a normal control. Diverging from the majority of transcriptomic studies that employ RNA-seq to identify inter-tissue differences, our approach reversed this rationale. We harnessed the sensitivity of RNA-seq and the representational power of autoencoders to conduct a comparative analysis aimed at identifying similarities—a common molecular signal that defines a shared identity between the peritumoral field and the tumor. Through this strategy within a three-arm design that included a truly normal tissue control, we successfully deconvoluted a 138-gene transcriptional signature defining the gastric field of cancerization. Our findings show that this signature is established early in the peritumoral field and persists in the established tumor, offering molecular evidence that carcinogenesis is a field process, not a focal event.

The analysis of representative genes within this signature offers a detailed portrait of this multifaceted process. Genes such as *ACSS1* (acetate metabolism) and *MSMO1* (cholesterol biosynthesis) directly point to the metabolic reprogramming required to sustain bioenergetic demand and cell proliferation. Simultaneously, the dysregulation of genes like *ITGB3* (cell adhesion), *MST1R* (motility), and *ADAMTS1* (matrix remodeling) indicates that the field is already being primed for an invasive phenotype. Furthermore, the activation of *XBP1* suggests that stress survival mechanisms are already operational. It is crucial to note that this same signature harbors the drivers *FAT4, MDM4*, and *NDRG1*.

Within this signature, the identification of the driver genes *FAT4, MDM4*, and *NDRG1* offers mechanistic insight into how this field is regulated. The overexpression of *FAT4*, a component of the Hippo tumor suppressor pathway, is an intriguing finding. Although the Hippo pathway is canonically suppressive, recent work in various cancers, including gastric cancer, suggests that the role of *FAT4* may be context-dependent, potentially acting oncogenically by disrupting cell polarity and cell-cell adhesion, crucial events for tumor initiation (Huang et al., 2022; Yang et al., 2012). Overexpression of *MDM4*, a potent p53 inhibitor, provides a clear mechanism for evading apoptosis and maintaining genomic instability. Inactivation of the p53 pathway is an almost universal event in cancer, and the amplification of *MDM4* is acknowledged as a key mechanism for accomplishing this in tumors that are wild-type for *TP53* (de Oliveira-Junior et al., 2021). The repression of *NDRG1*, in turn, aligns with its well-established role as a suppressor of the epithelial-mesenchymal transition (EMT) and metastasis (Li et al., 2021). Its early repression in the cancerization field may indicate that the cells are already being primed for a more invasive phenotype long before they become histologically malignant.

The positive correlation between *FAT4* and *MDM4*, exclusive to the peritumoral field, suggests a functional synergy required to establish the pre-neoplastic state. In contrast, the emergence of a negative correlation between *FAT4* and *NDRG1* only in the established tumor points to the activation of a distinct progression program, possibly related to invasion. This molecular “dance” among the drivers illustrates that carcinogenesis is not a linear process but an adaptive program that rewires its regulatory networks as it evolves.

From a translational perspective, the robust association of high *FAT4* and *MDM4* expression with an unfavorable prognosis elevates them to clinically relevant biomarkers. The validation of *MDM4*’s prognostic value in our local cohort of patients from Pará is particularly significant, as it demonstrates the clinical applicability of this finding across different populations.

The search for robust and validated biomarkers is one of the greatest challenges in gastric oncology, and our results contribute to this effort, aligning with other studies that seek to identify prognostic gene signatures.

Our study has limitations, including the relatively modest size of the validation cohort for the survival analysis. Nevertheless, it opens clear pathways for future investigations. The functional validation of the role of these three drivers in tumor initiation, using patient-derived gastric organoid models, is a logical and necessary next step.

In conclusion, our work redefines the gastric peritumoral tissue as a distinct, molecularly active pre-neoplastic field characterized by profound metabolic reprogramming. We identify *FAT4* and *MDM4* as drivers of this initial transition and as robust biomarkers of unfavorable prognosis, providing new insights into the earliest events of gastric carcinogenesis and opening new avenues for patient risk stratification.

## Supporting information

https://docs.google.com/document/d/1SNM9svID4TgCGJjRDOmWyWQWF_p6AiLyawju-Vhe0uc/edit?usp=sharing

## Acknowledgment

The authors would like to thank the Oncology Research Center, the Human and Medical Genetics Laboratory, and the Anatomical Pathology Laboratory at João de Barros Barreto University Hospital (HUJBB – UFPA) for their invaluable technical and laboratory support. Our gratitude also goes to the High-Performance Computing Center (CCAD) at the Federal University of Pará for access to the Apollo 2000 cluster, which was crucial for our analyses.

## Data availability statement

The original contributions presented in the study are included in the article. Further inquiries can be directed to the corresponding author.

