## Supplementary material for "Transcriptional Signatures of Field Cancerization in Gastric Cancer": https://docs.google.com/document/d/1SNM9svID4TgCGJjRDOmWyWQWF_p6AiLyawju-Vhe0uc/edit?usp=sharing

Figure S1. History of autoencoder training fo tumoral samples.

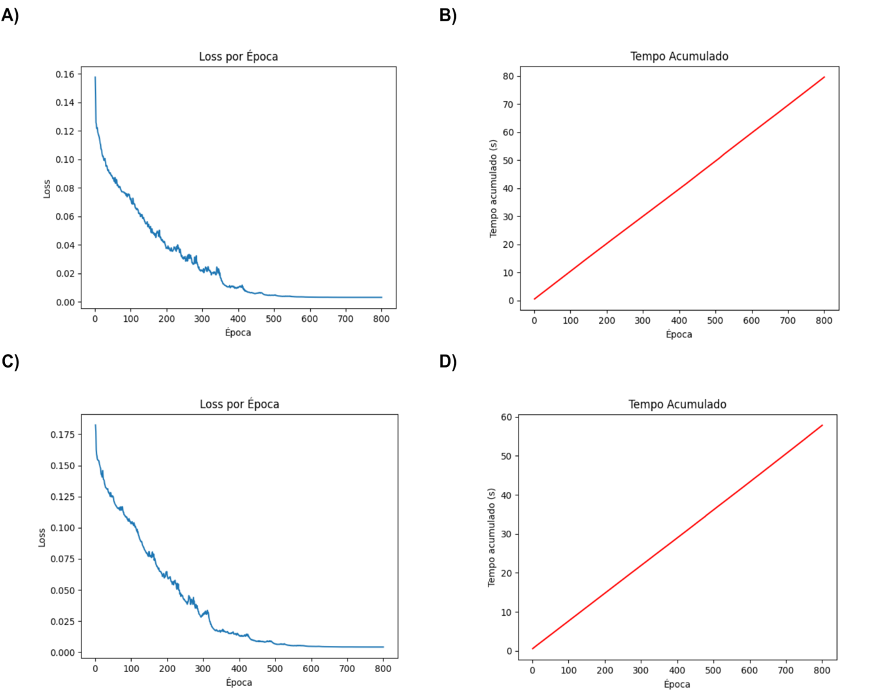

### Figure S2. History of autoencoder training for peritumoral samples.

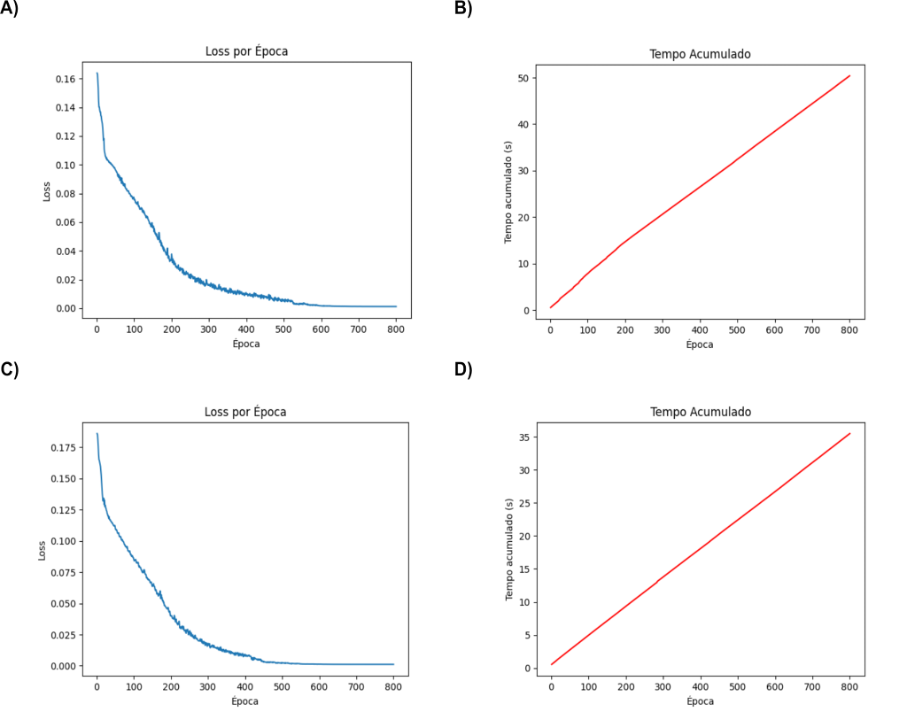

### **Materials and Methods**

**Study Cohorts and Sample Collection** Tumor and adjacent peritumoral tissues (collected >5 cm from the tumor margin) were obtained from 72 patients with gastric adenocarcinoma undergoing surgical resection at a reference hospital in the state of Pará, Brazil. All patients provided written informed consent. The study was approved by the local Research Ethics Committee. For a normal tissue reference, RNA-seq data from 68 non-cancerous gastric tissue samples were retrieved from the NCBI BioProject database under accession number PRJNA1054173.

**RNA Extraction, Library Preparation, and Sequencing** Total RNA was extracted from tissue samples using TRIzol® Reagent (Thermo Fisher Scientific) according to the manufacturer's protocol. RNA concentration and quality were assessed using a Qubit 4.0 Fluorometer (Thermo Fisher Scientific). Only samples with an RNA Integrity Number (RIN) > 7 were selected for downstream analysis. Purified RNA was stored at -80°C. RNA library preparation and sequencing were performed on a NextSeq® platform using the NextSeq® 500 MID Output V2 kit.

**RNA-Seq Data Processing and Normalization** Raw sequencing reads were demultiplexed and subjected to quality control and adapter trimming using Fastp (v0.23.2). Transcript quantification was performed using Salmon (v1.9.0) against the Gencode human reference transcriptome (v43). The tximport R package was used to summarize transcript-level quantifications to the gene level. Genes with fewer than 10 reads in at least 80% of samples were excluded from the analysis. The resulting gene count matrix was variance-stabilized using the regularized logarithm (rlog) transformation implemented in the DESeq2 R package. To ensure comparability across datasets, expression values for all samples (tumor, peritumoral, and normal) were subsequently scaled to a [0, 1] range using the MinMaxScaler function from the scikit-learn Python library, accessed via the reticulate R package.

**Deep Autoencoder Architecture and Training** A deep autoencoder was implemented using the Keras and TensorFlow libraries to learn a compressed, low-dimensional representation of the gene expression data. The final model architecture consisted of an input layer with neurons corresponding to the number of filtered genes, followed by three dense encoding layers (512, 256, and 64 neurons) with ReLU activation. The bottleneck (coding) layer contained 64 neurons. The decoder was composed of two dense layers (256 and 512 neurons) and an output layer with sigmoid activation to reconstruct the input. The model was compiled with the Adam optimizer and the Mean Absolute Error (MAE) loss function. Training was performed for 800 epochs with a batch size of 64. To prevent overfitting and optimize training, EarlyStopping (patience=20) and ReduceLROnPlateau (factor=0.2, patience=10) callbacks were employed. The model's reconstruction error was evaluated by calculating the MAE between the original and reconstructed data.

**Feature Importance and Gene Selection using SHAP** To ensure model interpretability, the latent space activations from the trained autoencoder were predicted using Random Forest models trained on the input gene expression data. The marginal contribution of each gene to the activation of each latent neuron was then quantified using SHAP (SHapley Additive exPlanations). For each neuron, the most relevant genes were selected by identifying the inflection point ('Elbow method') on the curve of rank-ordered positive SHAP values. This approach allowed for the data-driven selection of genes with the greatest impact on the model's learned representations.

**Statistical Pipeline for Field Signature Identification** A sequential four-step statistical pipeline was designed to isolate a stable gene signature characteristic of the peritumoral field and identify its key cancer drivers.

1. **Intersection of SHAP-derived Genes:** The gene lists identified by SHAP as relevant for the tumor and peritumoral autoencoder representations were intersected using the VennDiagram R package to create a core list of genes shared between both conditions.
2. **Selection for Expression Stability:** To identify genes that represent an early and stable alteration, we analyzed the expression of the shared genes between paired tumor and peritumoral samples from the same patient. A Wilcoxon signed-rank test was applied, and, contrary to conventional differential expression analysis, genes exhibiting high stability (i.e., no significant difference; p > 0.9) were selected for the next step.
3. **Differential Expression Analysis against Normal Tissue:** The list of stable genes was then compared to the normal tissue cohort. Differential expression analysis was performed using the DESeq2 R package, and only genes with a significant dysregulation (|log₂ Fold Change| > 1 and adjusted p-value < 0.05) were retained. This step ensured that the stable signature was specific to the carcinogenic field and absent in healthy tissue.
4. **Cross-referencing with Cancer Gene Databases:** Finally, to prioritize clinically relevant genes, the resulting list was cross-referenced with the COSMIC (Catalogue Of Somatic Mutations In Cancer) Cancer Gene Census database to identify established cancer driver genes.

**Visualization of Gene Expression Patterns** To visualize the expression patterns of the final 138-gene signature across all three tissue types (tumor, peritumoral, normal), a heatmap was generated using the Pheatmap R package. Prior to plotting, expression values for this gene set were converted to z-scores for row-wise scaling. For visualization, we computed Uniform Manifold Approximation and Projection (UMAP; R package umap) using Euclidean distances with default settings (*n_neighbors* = 15, *min_dist* = 0.1), and plotted samples by tissue class (healthy, adjacent, tumor) with normal-ellipse overlays. To quantify the signature per sample, we used single-sample GSEA (ssGSEA) from the GSVA package applied to the same VST matrix and custom 138-gene list, yielding one enrichment score per sample. Group differences (healthy vs. adjacent vs. tumor) were assessed on ssGSEA scores using the Kruskal–Wallis test followed by pairwise Wilcoxon tests with Benjamini–Hochberg correction

**Context-Dependent Co-expression Analysis** To investigate whether the regulatory relationships between the final driver genes (*FAT4*, *MDM4*, *NDRG1*) were altered during carcinogenesis, we performed co-expression analysis independently within the peritumoral and tumor sample groups. Pairwise Spearman's rank correlation coefficients (ρ) and corresponding p-values were calculated for the expression levels of these three genes in each context to identify statistically significant, tissue-specific correlations.

**Prognostic and Survival Analysis** The prognostic significance of the identified driver genes was evaluated through a two-tiered approach.

1. **External Cohort Analysis:** The association with overall survival (OS) was first assessed in large, publicly available gastric cancer cohorts using the Kaplan-Meier Plotter online database (kmplot.com). Patients were split by the median expression of each gene. Hazard ratios (HR), 95% confidence intervals, and log-rank p-values were retrieved.
2. **Internal Cohort Validation:** To validate these findings, an internal survival analysis was conducted on a subset of 46 patients from our Pará cohort for whom clinical follow-up data were available. Kaplan-Meier survival curves were generated using the survminer and survival R packages, and differences in OS based on median gene expression were assessed using the log-rank test. A p-value < 0.05 was considered statistically significant.
